# A bivalent EBV vaccine induces neutralizing antibodies that block B and epithelial cell infection and confer immunity in humanized mice

**DOI:** 10.1101/2022.01.18.476774

**Authors:** Chih-Jen Wei, Wei Bu, Laura A. Nguyen, Joseph D. Batchelor, JungHyun Kim, Stefania Pittaluga, James R. Fuller, Hanh Nguyen, Te-Hui Chou, Jeffrey I. Cohen, Gary J. Nabel

## Abstract

Epstein Barr virus (EBV) is the major cause of infectious mononucleosis and is associated with several human cancers. Despite its prevalence and major impact on human health, there are currently no specific vaccines or treatments. Four viral glycoproteins, gp 350 and gH/gL/gp42 mediate entry into the major sites of viral replication, B cells and epithelial cells. Here, we designed a nanoparticle vaccine displaying these proteins and show that it elicits potent neutralizing antibodies that protect against infection *in vivo*. Based on structural analyses, we designed single chain gH/gL and gH/gL/gp42 proteins that were each fused to bacterial ferritin to form a self-assembling nanoparticles. X-ray crystallographic analysis revealed that single chain gH/gL and gH/gL/gp42 adopted a similar conformation to the wild type proteins, and the protein spikes were observed by electron microscopy. Single chain gH/gL or gH/gL/gp42 nanoparticle vaccines were constructed to ensure product homogeneity needed for clinical development. These vaccines elicited neutralizing antibodies in mice, ferrets, and non-human primates that inhibited EBV entry into both B cells and epithelial cells. When mixed with a previously reported gp350 nanoparticle vaccine, gp350D_123_, no immune competition was observed. To confirm its efficacy in vivo, humanized mice were challenged with EBV after passive transfer of IgG from mice vaccinated with control, gH/gL/gp42+gp350D_123_ or gH/gL+gp350D_123_ nanoparticles. While all control animals (6/6) were infected, only one mouse in each vaccine group that received immune IgG had transient low level viremia (1/6). Furthermore, no EBV lymphomas were detected in immune animals in contrast to non-immune controls. This bivalent EBV nanoparticle vaccine represents a promising candidate to prevent EBV infection and EBV-related malignancies in humans.

**One sentence summary:** A bivalent gp350 and gH/gL/gp42 nanoparticle vaccine elicits neutralizing antibodies that protect against EBV infection and EBV lymphoma *in vivo*.

## INTRODUCTION

Over 95% of adults worldwide are infected with Epstein-Barr virus (EBV), the primary agent for infectious mononucleosis (IM) (*1*). EBV was discovered in the 1960s and was the first human virus associated with cancer; EBV has since been associated with malignancies such as nasopharyngeal carcinoma, Hodgkin’s lymphoma, non-Hodgkin’s lymphoma, Burkitt’s lymphoma, NK/T cell lymphomas, peripheral T-cell lymphomas and gastric cancer (*1, 2*). Each year, more than 200,000 cases of cancer are associated with EBV infection, resulting in ∼140,000 deaths (*2*). EBV is also the main cause of lymphoproliferative disease in patients with immunodeficiencies. Nearly all post-transplant lymphoproliferative disorder (PTLD) in the first year after is caused by EBV (*3*).

There is currently no therapy to effectively treat EBV infection, and there is no vaccine to prevent EBV infection. Prior vaccine development attempts mainly focused on one of the viral envelope glycoproteins gp350 as it is the most abundant surface protein and is the major target of neutralizing antibodies (*4*). gp350 mediates viral entry to B cells by engaging complement receptor 2 (CR2/CD21) (*5*). Other viral surface glycoproteins, namely gH, gL, gB, gp42 and BMRF2 also play a role in EBV infection and are also targets of neutralizing antibodies. gp42 binds to human leukocyte antigen (HLA) class II and together with gH/gL heterodimer and gB forms a complex that promotes EBV entry to B cells (*6*). Infection of EBV to epithelial cells is initiated by the engagement of BMRF2 and gH/gL complex with integrin receptors and ephrin receptor A2 (*7–9*). A prototype gp350 vaccine reduced the incidence of IM by 78%, but did not prevent infection in a phase II clinical trial (*10*). Other gp350-based vaccines have also shown protective efficacy in relevant nonhuman primate models (*7, 11, 12*). Recombinant gH/gL complex or gB have also been shown to induce neutralizing antibody responses in rabbits (*13*). We have previously shown that a nanoparticle (NP)-based gp350 vaccine elicited protective immunity (*14*), and gH/gL and gH/gL/g42 NP vaccines induced potent neutralizing antibodies that inhibit EBV entry in both B cells and epithelial cells (*15*). In this study, we optimized the consistency of gH/gL and gH/gL/gp42 that could reduce the heterogeneity of these heteromeric proteins by generating a single chain polypeptide based on structural biology. This approach not only preserved the presentation of these antigens but also ensured that the gH/gL and gp42 heteromers were uniformly assembled to improve product consistency required for clinical grade vaccines. Here, we have evaluated the structure, immunogenicity and protection of single chain gH/gL-NP or single chain gH/gL/gp42-NP together with gp350-NP in relevant animal models.

## RESULTS

### Design and characterization of single chain gH/gL and gH/gL/gp42 nanoparticles

When co-transfection of plasmids is used to generate multimeric complexes (*15*), there is potential inconsistency in the product if the appropriate stoichiometry is not achieved. To address this concern, we generated single chain gH/gL or gH/gL/gp42 complexes using structural data to fuse the ectodomains with flexible linkers (Fig. 1A). Fusion to specific sites on ferritin (*14*) facilitated the formation of self-assembling nanoparticles (NP). The gH/gL and gH/gL/gp42 fusion proteins migrated as a single band on SDS-PAGE gel (Fig. S1). Crystal structures of single chain gH/gL and single chain gH/gL/gp42 were determined (Fig. 1B; Supplemental Table 1). Each structure superimposed on previously published heterodimeric gH/gL (PDB 3PHF) and heterotrimeric gH/gL/gp42 (PDB 5T1D) complex structures, respectively, demonstrating that single chain gH/gL and single chain gH/gL/gp42 adopt native conformations resembling the wild-type complexes (Fig. 1B). Both gH/gL-NP and gH/gL/gp42-NP could be purified by ion exchange and size-exclusion chromatography (Fig. 1C, left), and dynamic light scattering analysis documented the expected particle radius of 20.7 nm and 24.8 nm for single chain gH/gL-NP and single chain gH/gL/gp42-NP, respectively (Fig 1C, right). The single chain gH/gL-NP and single chain gH/gL/gp42-NP were also visualized by transmission electron microscopy, showing visible spikes protruding from the ferritin core, consistent with the expected structure and stoichiometry (Fig. 1D, left and right panels respectively).

**Figure 1.**
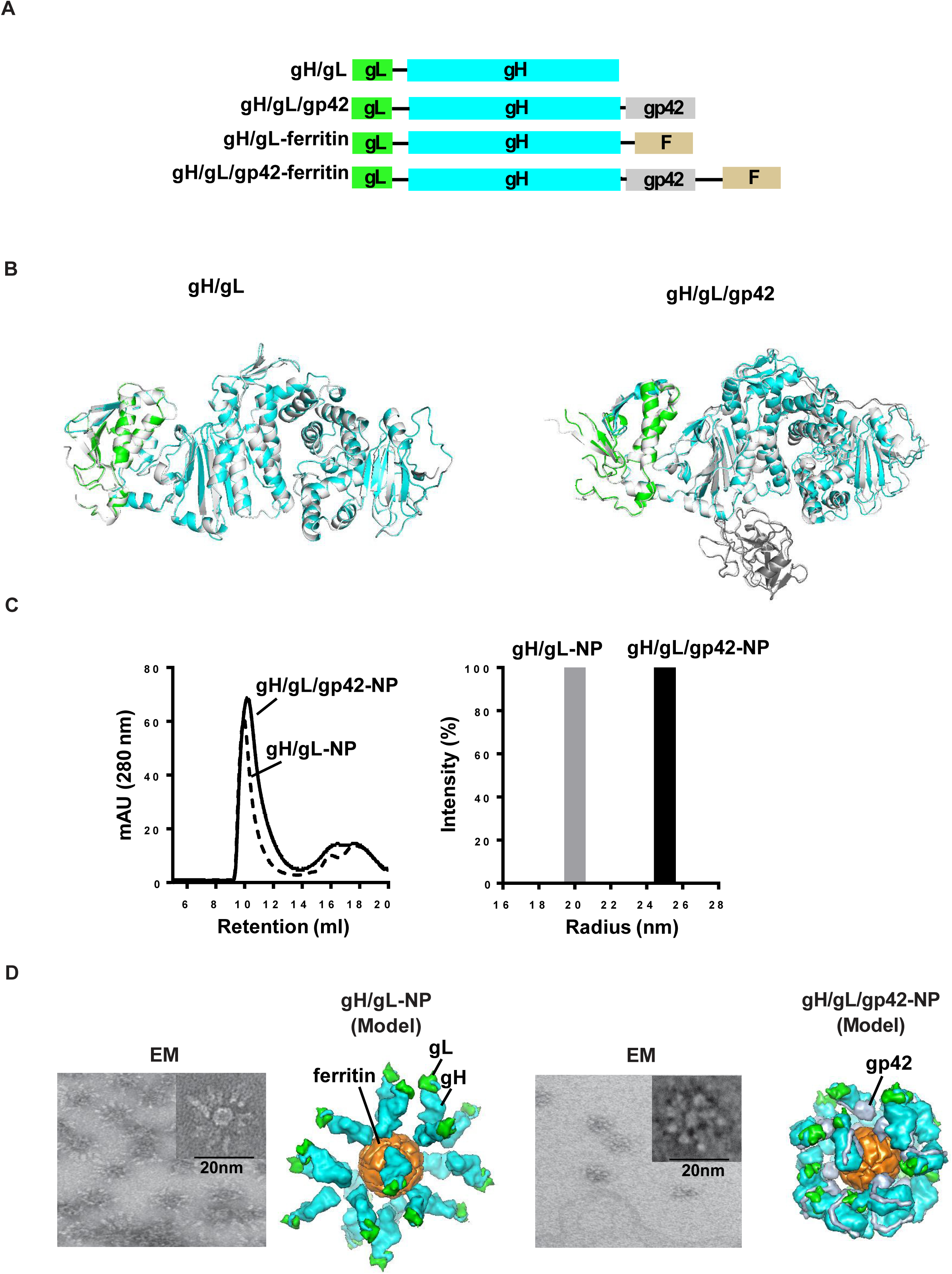
Structure-based design of single chain gH/gL and single chain gH/gL/gp42 nanoparticles. **(A)** A schematic representation of the single chain gH/gL, single chain gH/gL/gp42, single chain gH/gL-NP, and single chain gH/gL/gp42-NP. EBV gL (green) is fused to the N terminus of gH (cyan) via a flexible amino acid linker (indicated by the black line between gL and gH). EBV gp42 (gray) is fused to the C-terminus of gH. Single chain gH/gL-NP or single chain gH/gL/gp42-NP constructs show the gH/gL or gH/gL/gp42 fused to *H. pylori* ferritin (represented by the letter “F” in beige) by a flexible amino acid linker (line), respectively. **(B)** Left: The crystal structure of the single chain gH/gL was resolved at 5.5Å (gL in green and gH in cyan) with superposition of the previously solved crystal structure of gH/gL complex (white, PDB: 3PHF) (RMS = 0.33). Right: The crystal structure of the single chain gH/gL/gp42 was resolved at 2.9Å and superposition with the previously solved heterotrimer gH/gL/gp42 complex crystal structure (white, PDB: 5T1D) (RMS value = 0.96). **(C)** Left: Size exclusion chromatography (SEC) elution profiles of single chain gH/gL-NP and single chain gH/gL/gp42-NP. Right: Size of single chain gH/gL-NP and single chain gH/gL/gp42-NP determined by DLS. **(D)** Negative stain EM image of single chain gH/gL-NP and single chain gH/gL/gp42-NP. A close-up image of the nanoparticle is displayed at the upper right corner. A structural model of the single chain gH/gL-NP or single chain gH/gL/gp42-NP is shown on the right of the EM image (gH: cyan; gL: green; gp42: gray; ferritin: orange). The surface density is a model built from crystal structures solved in panel B and ferritin core from PDB 3BVE (DOI:10.2210/pdb3bve/pdb) using Chimera (*40*) and is not reconstructed from EM.

### Immunogenicity of single chain gH/gL-NP or gH/gL/gp42-NP in mice

The immunogenicity of single chain gH/gL-NP was first evaluated in mice with or without AF03, a squalene-based oil-in-water emulsion adjuvant previously used in a pandemic influenza vaccine (*16*). The anti-gH/gL antibody titer was significantly higher in the adjuvanted group after each immunization (Supplemental Fig. 2, p<0.0001). All subsequent animal studies were therefore carried out in the presence of AF03 adjuvant. Single chain gH/gL-NP elicited a robust antibody response against the gH/gL complex and when used together with the previously described gp350D_123_-NP (*14*), no immune competition was seen (Supplemental Fig. 3). Similar results were observed when gp42 was included to generate a single chain gH/gL/gp42-NP that induced antibodies against both gH/gL and gp42 after immunization (Supplemental Fig. 4A). Again, no immune competition was observed when mice were given a bivalent single chain-gH/gL/gp42-NP and gp350D_123_-NP vaccine as the antibodies titers against each individual antigen remained at the similar levels (Supplemental Fig. 4B). The neutralizing activity of sera from animals immunized with single chain gH/gL-NP and single chain gH/gL/gp42-NP were evaluated in both B cells and epithelial cells (Fig 2). Single chain gH/gL-NP, gH/gL/gp42-NP and gp350D_123_-NP all elicited potent neutralizing antibodies that blocked virus entry to B cells, and the neutralizing IC_50_ titers remained at similar or higher levels when single chain gH/gL-NP or single chain gH/gL/gp42-NP was mixed with gp350D_123_-NP in a bivalent vaccine (Fig. 2A, 2B, left, p<0.05 compared to control). Virus neutralization in epithelial cells was also evident in mice that received single chain gH/gL-NP and single chain gH/gL/gp42-NP while gp350D_123_-NP antiserum had minimum effect in blocking EBV infection in epithelial cells, consistent with our previous report (*15*) (Fig. 2A, right, p<0.05 compared to control). Again, bivalent single chain gH/gL-NP+gp350D_123_-NP or single chain gH/gL/gp42-NP+gp350D_123_-NP induced similar neutralizing antibody titers as the monovalent single chain gH/gL-NP and single chain gH/gL/gp42-NP, respectively (Fig. 2A, 2B, right, p<0.05 compared to control).

**Figure 2.**
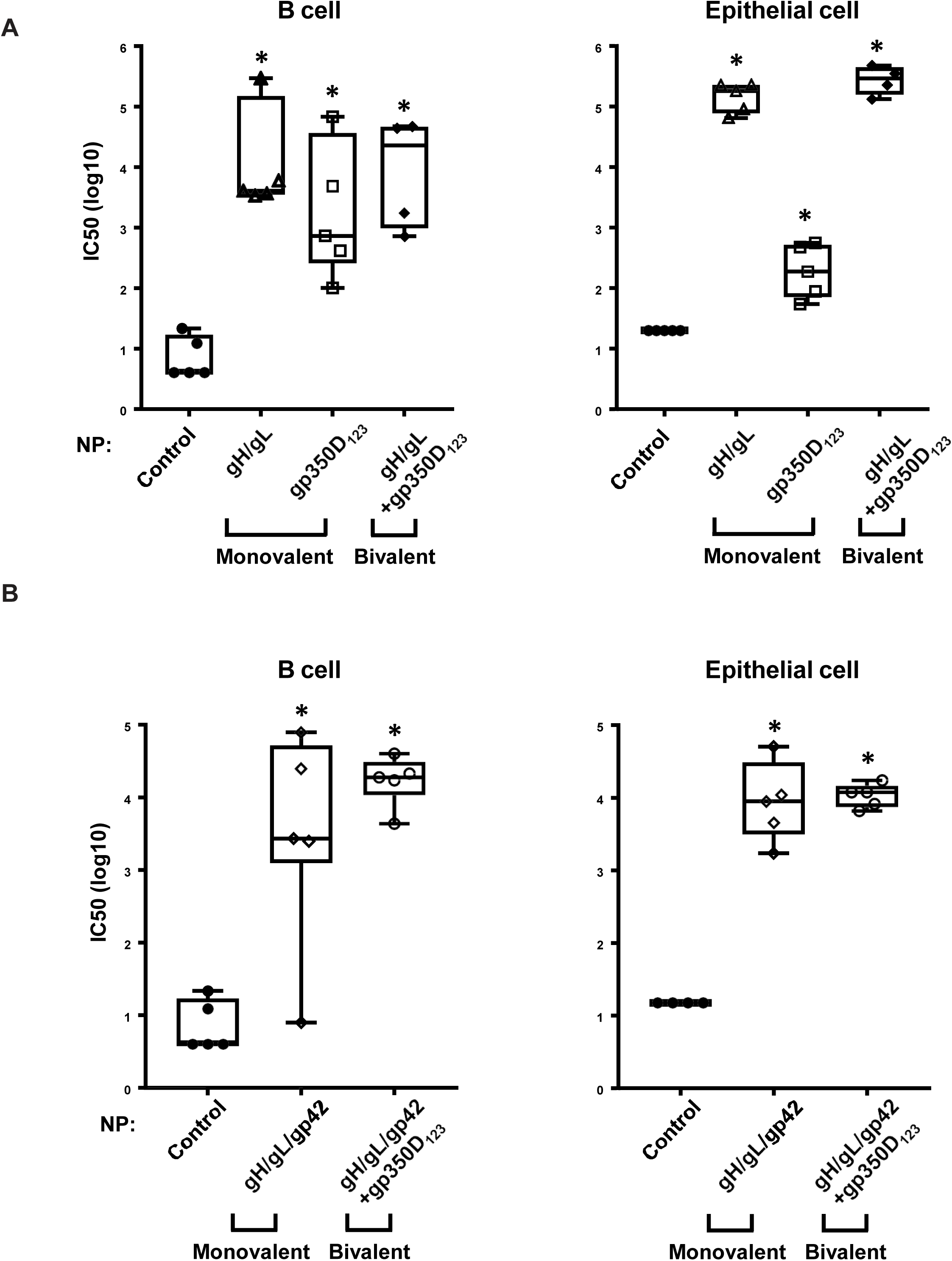
Neutralization responses induced by single chain gH/gL-NP or single chain gH/gL/gp42-NP alone or in combination with gp350D_123_-NP in mice. BALB/C mice (n=5/group) were immunized intramuscularly in the presence of AF03 adjuvant at weeks 0 and 3 with 1µg of **(A)** monovalent single chain gH/gL-NP, gp350D_123_ or bivalent gH/gL-NP+gp350D_123_-NP or **(B)** monovalent single chain gH/gL/gp42-NP or bivalent gH/gL/gp42-NP+gp350D_123_-NP. Control is pre-immune sera. Neutralization antibody titers from immune sera collected 2 weeks after the 2^nd^ injection was determined in Raji B cells and SVK CR2 epithelial cells. The IC_50_ indicated the log titer that resulted in 50% inhibition of EBV entry into target cells. The data are shown as box-and-whiskers plots (box indicates lower and upper quartiles with line at median, and whiskers span minimum and maximum data points; *p<0.05 compared to control).

### Bivalent single chain gH/gL-NP+gp350D_123_-NP or single chain gH/gL/gp42-NP+gp350D_123_-NP elicited neutralizing antibodies in ferrets and non-human primates

We next evaluated the immunogenicity of bivalent single chain gH/gL-NP+gp350D_123_-NP and single chain gH/gL/gp42-NP+gp350D_123_-NP in EBV-naïve ferrets. Non-immune ferrets were immunized with 2 doses of single chain gH/gL-NP+gp350D_123_-NP or single chain gH/gL/gp42-NP+gp350D_123_-NP at weeks 0 and 4. Immune sera from ferrets receiving single chain gH/gL/gp42-NP+gp350D_123_-NP elicited more potent neutralizing antibodies that inhibit virus entry to B cells than animals immunized with single chain gH/gL-NP+gp350D_123_-NP (Fig 3A, left, p<0.05 compared to pre-immune sera). In epithelial cells, both single chain gH/gL-NP+gp350D_123_-NP and single chain gH/gL/gp42-NP+gp350D_123_-NP anti-sera showed high titers of antibodies that neutralized EBV infection (Fig 3A, right, p<0.05 compared to pre-immune sera). These findings were confirmed by measuring ELISA binding of antibodies to each component elicited by each bivalent vaccine (Supplemental Fig. 5, **p<0.0001 or *p<0.05 respectively compared to pre-immune sera). Background reactivity observed in pre-immune ferret sera likely represented non-specific binding of the secondary anti-ferret antibody as it was not observed in the neutralization assay. Despite this background, antibody levels increased 9-400-fold after immunization.

**Figure 3.**
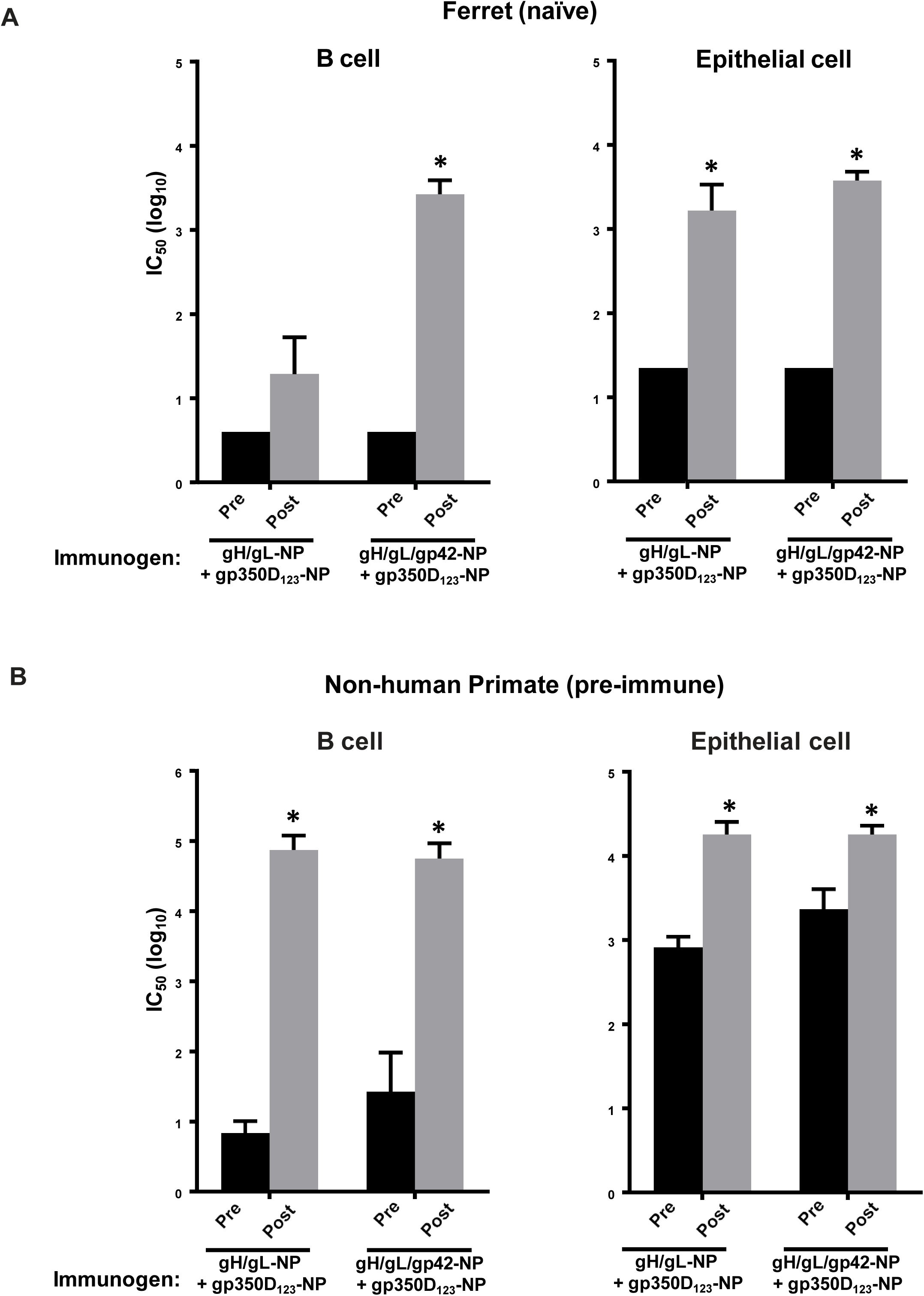
Immunogenicity of single chain gH/gL-NP+gp350D_123_-NP or single chain gH/gL/gp42-NP alone or in combination with gp350D_123_-NP in ferrets and NHP. **(A)** Ferrets were immunized intramuscularly at weeks 0 and 4 with either 15 μg single chain gH/gL-NP + 15 μg gp350D_123_-NP or 15 μg single chain gH/gL/gp42-NP + 15 μg gp350D_123_-NP bivalent vaccines. Sera were collected 2 weeks after immunization and were assayed for neutralizing activity in both Raji B cells (left) and 293 epithelial cells (right). Means and standard deviations are shown (* p< 0.05 compared to pre-immune sera). No neutralizing activity was detected from the pre-immune sera and levels were at the limit of detection in the graphs. **(B)** Rhesus macaques (n=4/group) were vaccinated with the bivalent vaccine composed of 25µg gH/gL-NP + 25µg gp350D_123_-NP or 25µg gH/gL/gp42-NP + 25µg gp350D_123_-NP at weeks 0, 4, and 10. AF03 was used as adjuvant. Immune sera were collected 2 weeks after the 3^rd^ injection and neutralizing antibody titers were determined in both Raji B cells and SVK CR2 epithelial cells. Means and standard deviations are shown (* p< 0.05 compared to pre-immune sera).

A high percentage of rhesus macaques are infected naturally during infancy with rhesus lymphocryptovirus (rhLCV), a herpesvirus closely related and immunologically cross-reactive with EBV (*17*). To determine whether vaccination could boost pre-existing immune responses, we immunized rhesus macaques (*Macaca mulatta*) with single chain gH/gL-NP+gp350D_123_-NP or single chain gH/gL/gp42-NP+gp350D_123_-NP to model immune responses in humans. After the third immunization, serum neutralization titers in animals receiving single chain gH/gL-NP+gp350D_123_-NP or single chain gH/gL/gp42-NP+gp350D_123_-NP were substantially elevated compared to the pre-immune sera (Fig 3B, left; 10,000-fold for single chain gH/gL-NP+gp350D_123_-NP and >2,000-fold for single chain gH/gL/gp42-NP+gp350D_123_-NP, * p< 0.05 compared to pre-immune sera). As expected, substantial reactivity was observed in pre-immune sera (Supplemental Fig. 6, top and bottom left), likely due to the high amino acid sequence identity between EBV and rhLCV gH (85.4%), gL (81.8%) and gp42 (88%) (*17*). Similar to the pre-existing anti-gH/gL and anti-gH/gL/gp42 binding antibodies, neutralizing activity was observed in epithelial cells with the pre-immune monkey sera. Nonetheless, the neutralizing antibody titers increased by more than 20- and ∼7-fold after the third immunization with single chain gH/gL-NP+gp350D_123_-NP and single chain gH/gL/gp42-NP+gp350D_123_-NP, respectively (Fig. 3B, right) and the activity was maintained for at least 12 weeks (data not shown). These findings were confirmed by measuring ELISA binding titers to gH/gL and gH/gL/gp42. In contrast, little reactivity to gp350D_123_ was observed in pre-immune sera because the homology between EBV and rhLCV gp350 is much lower (49%) (Supplemental Fig 6, top and bottom right). gp350 is important for EBV attachment to B cells before the gH/gL/gp42 complex initiates membrane fusion and is likely the reason why there was minimal difference in neutralization between B and epithelial cells. Regardless of prior immunity, anti-gH/gL, anti-gH/gL/gp42 and anti-gp350 antibody titers all increased after immunization (Supplemental Fig 6, **p<0.0001 or *p<0.01 respectively compared to pre-immune sera). These data confirmed the immunogenicity of bivalent single chain gH/gL-NP+gp350D_123_-NP and single chain gH/gL/gp42-NP+gp350D_123_-NP vaccines and indicate that neutralizing antibodies were induced by these vaccines in nonhuman primates. Although there is variation in antibody titers elicited by the bivalent vaccine in different models, our data clearly demonstrate that neutralizing antibodies could be readily elicited by the bivalent vaccines described in both naïve and EBV-immune animal models.

### Bivalent single chain gH/gL-NP+gp350D_123_-NP or single chain gH/gL/gp42-NP+gp350D_123_-NP immune sera protected humanized mice from EBV infection and lymphoma

To assess the protective efficacy of single chain gH/gL-NP+gp350D_123_-NP and single chain gH/gL/gp42-NP+gp350D_123_-NP vaccines, we performed an EBV challenge study using a humanized mouse model. Engraftment of human CD34+ hematopoietic stem cells into NOD-scid IL2rg^-/-^ mice (CD34+ huNSG) allows for the reconstitution of human immune system components and these mice can be infected with EBV (*18–21*). Purified IgG from naïve (control), single chain gH/gL-NP+gp350D_123_-NP or single chain gH/gL/gp42-NP+gp350D_123_-NP immunized BALB/c mice were passively transferred to three groups of CD34+ huNSG mice on days -1, 0 and +1 and mice were challenged intravenously with EBV on day 0. All animals in the control group had viremia while only one animal each receiving IgG from single chain gH/gL-NP+gp350D_123_-NP or single chain gH/gL/gp42-NP+gp350D_123_-NP immune animals had low level transient viremia on one day, further demonstrating the protective efficacy conferred by single chain gH/gL-NP+gp350D_123_-NP and single chain gH/gL/gp42-NP+gp350D_123_-NP vaccines (Fig. 4A, p<0.05 at week 5 and week 9). In situ hybridization for EBV encoded RNA 1 (EBER1) showed viral RNA in tissue of 4 of 6 animals receiving control IgG, while tissues from all animals in the other two groups were entirely negative for EBER1 (Fig. 4B, p<0.005). Three of the six animals that received control IgG had EBV positive B cell lymphomas, while none of the animals that received IgG from either single chain gH/gL-NP+gp350D_123_-NP or single chain gH/gL/gp42-NP+gp350D_123_-NP immune animals developed lymphoma (Fig. 5 and Supplemental Fig. 7, p<0.05). We also determined the levels of anti-gH/gL, anti-gH/gL/gp42 and anti-gp350 in animals 1 week after challenge using an immunoprecipitation assay, and the antibody levels against gp350, gH/gL and gp42 were comparable to those from animals immunized with 2 doses of EBV vaccines in previous studies, ranging from 10^5^ to 10^6^ relative light units (Supplemental Fig. 8) (*14, 15*).

**Figure 4.**
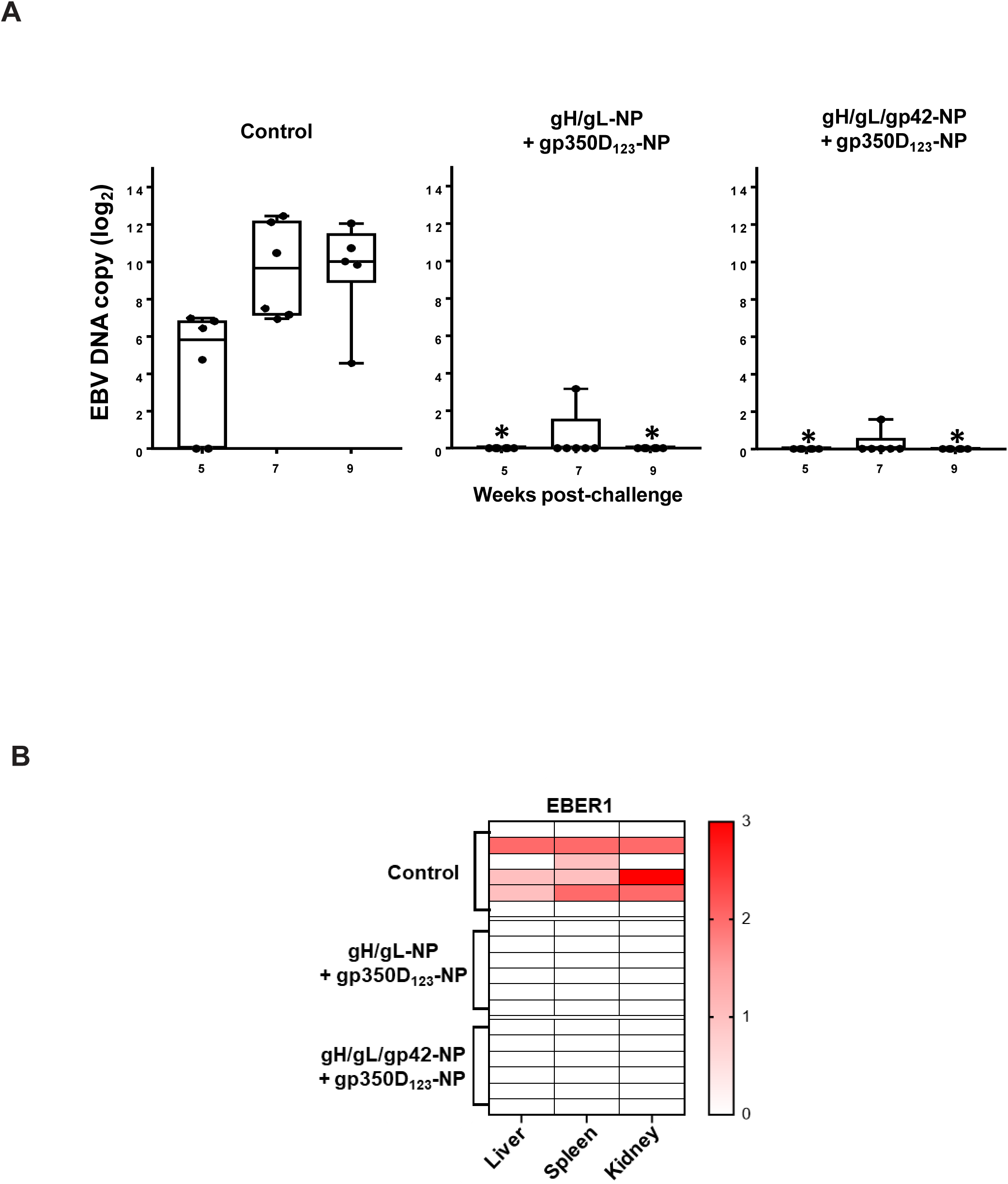
Immune protection by passive transfer of bivalent vaccine sera against EBV infection in humanized NSG mice. Humanized NSG mice (n=6/group) were injected IgG (20µg/g of mouse body weight) purified from naïve (control), single chain gH/gL-NP+ gp350D_123_-NP or single chain gH/gL/gp42+gp350D_123_-NP immunized BALB/C mice. Passive transfer of IgG was delivered intraperitoneally on day -1, 0 and 1 and EBV challenge was performed intravenously on day 0. **(A)** Viremia from each group was measured at weeks 5 7, and 9 post challenge. Medians with 25% and 75% percentiles are shown (* p< 0.05 compared to control at the same week). **(B)** Heatmap showing EBV encoded RNA 1 (EBER1) positivity of tissues (graded 0 to 3) from mice receiving IgG from naïve, single chain gH/gL-NP+ gp350D_123_-NP, or single chain gH/gL/gp42+gp350D_123_-NP immunized BALB/C mice after challenge with EBV. A score of 0 indicates no EBER1 staining while a score of 3 indicates marked infiltration of tissues by EBER1-positive cells.

**Figure 5.**
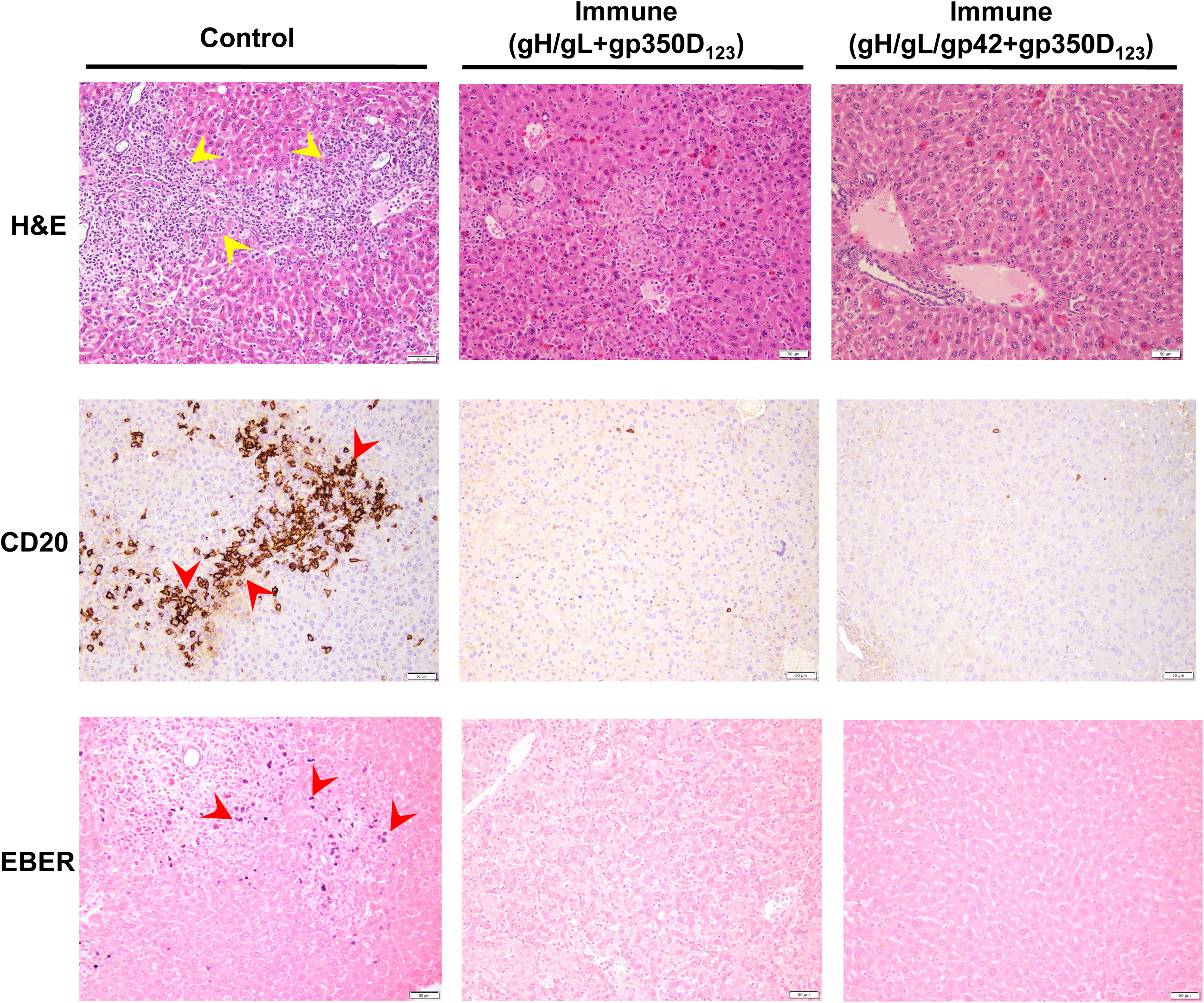
Protection against EBV lymphoma *in vivo*. Pathologic and immunohistochemical analysis of the liver from representative mice receiving IgG from non-immune (Control, left) or vaccinated mice (Immune, middle and right) after challenge with EBV. Tissues were collected 27 weeks after challenge and stained with hematoxylin and eosin (H&E, yellow arrows indicate representative region with lymphoma) or anti-CD20 antibody (brown staining, red arrows), or in situ hybridization was performed with a probe to EBER1 (purple staining, red arrows). CD20 and EBER staining are apparent in control, but not in any of the tissues receiving IgG from vaccinated mice. No EBV-positive B cell lymphomas were observed in the latter animals.

## DISCUSSION

An effective vaccine could reduce the burden of a variety of diseases associated with EBV infection, including infectious mononucleosis and a wide range of B cell and epithelial cell cancers. A previous phase 2 trial with an EBV gp350 vaccine reduced rate of infectious mononucleosis, but failed to induce sterilizing immunity in clinic (*10*), suggesting that additional immunogens that target viral entry to epithelial cells may be required for a successful prophylactic vaccine (*15, 22*). While EBV gp350 is important for attachment of the virus to B cells, it is not required for infection *in vitro*; in contrast, EBV gH, gL and gp42 are all essential for infection and EBV fusion to host cells. Here, we rationally designed vaccine candidates based on knowledge of their structural biology that allow expression of gH/gL and gH/gL/gp42 in a nanoparticle as a single polypeptide. This approach ensures the proper formation of 1:1 heterodimers for gH/gL and 1:1:1 heterotrimers for gH/gL/gp42, respectively. These single chain recombinant proteins can be easily purified and the conformation of the gH/gL and gH/gL/gp42 nanoparticles remains intact compared with nanoparticles produced from combinations of polypeptides as determined by x-ray crystallography. The ability to express the gH/gL or gH/gL/gp42 nanoparticles as single polyproteins reduces the number of components required to generate the vaccines and enables greater control of product homogeneity that facilitates scaled manufacturing.

Immunization of mice with single chain gH/gL-NP or single chain gH/gL/gp42-NP induced high titers of antibodies that neutralized EBV entry to B cells and epithelial cells. Addition of a structurally optimized, truncated gp350, gp350D_123_-NP (*14*), to single chain gH/gL-NP or single chain gH/gL/gp42-NP stimulated effective gp350 directed Abs without reducing gH/gL or gH/gL/gp42 responses as shown previously for gp350D_123_-NP and gH/gL-NP or gH/gL/gp42-NP (*15*), suggesting that a bivalent vaccine formulation with both gp350 and gH/gL/gp42 components would provide improved coverage against the virus on different cell types. These neutralizing antibody responses were observed in ferrets, an EBV-naïve mammalian species. Importantly, in monkeys with pre-existing cross-reactive immunity to EBV due to prior infection by rhLCV, substantially higher titers of neutralizing antibodies were induced by immunization with these bivalent vaccines. In mice and NHPs, both single chain gH/gL-NP + gp350D_123_-NP and gH/gL/gp42-NP + gp350D_123_-NP bivalent vaccines induced neutralizing antibodies that inhibited EBV entry in B cells and epithelial cells. The neutralizing antibody levels (IC_50_ titers) elicited by the bivalent vaccines were similar to those seen previously with combined gH/gL/gp42 + gp350 nanoparticle vaccines in other formats (*15*). Importantly, we show here that this immunity also protects against EBV infection and development of EBV lymphoma in vivo.

Evaluation of the protective efficacy of EBV vaccines is challenging as humans are the only natural reservoir for EBV. There are limitations in the animal models to evaluate EBV vaccine efficacy because EBV does not naturally infect rodents (*1*). In contrast, rhesus macaques are naturally infected almost universally by rhesus lymphocryptovirus (rhLCV) which is homologous to EBV; each of the rhLCV glycoproteins has an ortholog in EBV and antibodies to the rhLCV glycoproteins complicate EBV challenge studies. Furthermore, it is extremely difficult to obtain sufficient rhLCV seronegative NHPs for vaccination and challenge studies. Although vaccinated common marmosets have reduced EBV DNA in buccal fluid after EBV challenge (*23*), reduced shedding in oral fluids is not a useful test for efficacy of a vaccine. In addition, these animals are naturally infected with a marmoset homolog of EBV (*Callitrichine herpesvirus* 3) which has orthologs of EBV gH/gL/gp42 and gp350; thus, most common marmosets have antibodies to these glycoproteins that confound challenge studies. Finally, while rabbits can be infected with EBV, it requires non-physiologic, extremely high doses (20-80 million copies of EBV) of virus to infect them (*24*).

Several laboratories have modeled EBV infection in humanized mice engrafted with CD34+ hematopoietic progenitor cells isolated from umbilical cord blood (CD34+ huNSG) (*28*) and have shown that these animals become viremic after infection (*20*). Here we used this model to validate protection against EBV viremia using our bivalent EBV vaccines by passive transfer of immune IgG from vaccinated mice. Nearly all CD34+ huNSG mice that received purified IgG from bivalent vaccinated animals displayed undetectable viremia after EBV challenge; in contrast, 10^2^-10^3^ copies of EBV DNA/10 μl of blood were present in animals that received control IgG. Successful protection was conferred by passive transfer of vaccine-induced antibodies. Because it is technically not possible to perform active immunization in a relevant model of infection, we showed here that transfer of IgG obtained from serum alone conferred protection, demonstrating both the efficacy and mechanism of immune protection at the same time. There is ample precedent for the use of passive transfer to demonstrate vaccine-induced immune protection for other viruses (*25–27*). Similar to our previous report (*15*), both single chain gH/gL-NP and gH/gL/gp42-NP induced robust antibody responses that neutralized EBV entry to B cells and epithelial cells. The rationale to include gp42 is to mimic the natural heterotrimeric viral structure of the gH/gL/gp42 complex. This complex has been implicated in mediating B cell neutralization (*4*) and will be the lead candidate moving forward. Our approach ensures correct complex stoichiometry, simplifies protein production, minimizes heterogeneity, improves immunogen stability, and reduces manufacturing costs. Taken together, these data suggest that the single chain gH/gL/gp42 and gp350 bivalent vaccine represents an efficient, scalable candidate vaccine that is likely to limit viremia after EBV infection, thereby reducing infectious mononucleosis and possibly EBV associated cancers.

## MATERIALS AND METHODS

### Vector Construction

The EBV glycoproteins gH, gL and gp42 amino acid sequences were obtained from NCBI GenPept with the following accession numbers: gH (Q3KSQ3.1), gL(P03212.1) and gp42 (P0C6Z5.1). Through structural modeling, the glycoproteins were fused via a flexible amino acid linker to make a single chain gH/gL heterodimer or gH/gL/gp42 heterotrimer recombinant protein (Supplementary Table 2). A 6-histidine tag with a thrombin cleavage sequence was placed at the C-terminus of the single chain recombinant protein for affinity purification purposes. The EBV nanoparticle glycoproteins were generated by fusing the singe-chain glycoproteins to the N-terminus of *Helicobacter pylori*- ferritin (*14*).

### Recombinant protein expression and purification

Expi293F were transiently transfected using ExpiFectamine 293 reagent at a cell density of 2 x 10^6^ cells/ml with the gH/gL, gH/gL/gp42, gH/gL nanoparticle, gH/gL/gp42 nanoparticle, or gp350D_123_ nanoparticle expression vector (Life Technologies). After 5 days of expression the supernatants were harvested for purification. The gH/gL or gH/gL/gp42 constructs were affinity tagged with a hexahistidine tag and was purified via Ni Sepharose 6 Fast Flow histidine-tagged protein purification resin (GE Healthsciences). Nickel column eluate was concentrated to approximately 10 mg/mL using 10 kDa cutoff centrifugal filters. This material was then subjected to gel filtration chromatography using a HiLoad 16/600 Superdex 200pg column that had been equilibrated in TBS buffer (20 mM Tris pH 7.4, 150 mM NaCl). Peak fractions containing pure fusion protein (as judged by SDS-PAGE) were pooled and concentrated back to approximately 10 mg/mL. This sample was then deglycosylated by adding PNGase F enzyme at a ratio of 5 units PNGase F per µg of fusion protein and incubated at room temperature for 72 hours. PNGase F was then removed by gel filtration over a Superose 6 10/300 GL column equilibrated in TBS, pooling non-void fractions containing the fusion protein, but not PNGase F. These fractions were concentrated to 7.5 mg/mL and stored at 4°C. The nanoparticles were purified via ion exchange chromatography (20 mM Tris-HCl pH 7.5, 50 mM NaCl), followed by Superose 6 10/300gL size exclusion chromatography filtration column in PBS (GE Healthsciences). SDS-PAGE and western blots were performed to detect the presence of the nanoparticles (Biorad). Endotoxin analyses ensured that all vaccine doses contained <0.1 EU per mouse.

### Crystallization and cryoprotection

Crystallization was carried out by sitting drop vapor diffusion at 18°C against a solution of 0.1 M Bis-Tris pH 5.5, 0.375 M ammonium sulfate, and 19.5% PEG 3350 for single-chain gH/gL/gp42 and 1M LiCl, 10% PEG 6k, 0.1M Na_3_Citrate pH 5.0 for single-chain gH/gL. Drops (200 nL total volume) were set up at a 1:1 ratio of protein stock (7.5 mg/mL) and crystallization solutions. Crystals were cryo-protected by transfer into a fresh drop of the crystallization solution supplemented to 25% glycerol and incubated for 10 s immediately prior to freezing in liquid N_2_. X-ray diffraction data for single chain gH/gL were collected at the Advanced Photon Source beamline LS-CAT 21-ID-D and on an EigerX 9M Detector (wavelength 1.1 Å). X-ray diffraction data for single chain gH/gL/gp42 were collected at Diamond Light Source beamline i24 on a Pilatus 3 6M detector (wavelength 0.9686 Å). Both datasets were indexed, integrated and scaled using XDS (*29, 30*). Initial phases were obtained by molecular replacement with Phaser (*31, 32*) using the 3PHF structure for single chain gH/gL and the gH/gL/gp42 domains of 5W0K for single chain gH/gL/gp42. Structures were modeled and refined using the programs COOT (*33*) and PHENIX (*34*).

### Negative stain transmission electron microscopy

1 mg/mL of nanoparticle samples were sent to the Harvard Medical School Electron Microscopy Facility for negative stain transmission electron microscopy. The samples were stained with 0.75% uranyl formate and a TecnaiG^2^ Spirit BioTWIN microscope was used to image the grids. The images were recorded with an AMT 2k charge-coupled device camera.

### Immunization

Animal experiments were carried out in accordance with all federal regulations and were approved by the Sanofi Institutional Animal Care and Use Committee in fully AAALAC accredited facilities. Six- to eight-week old female BALB/c mice (Sanofi in house) were immunized (n=5) intramuscularly with purified proteins either in the absence or presence of Sanofi Pasteur AF03 adjuvant at 50% (v/v) formulation. 1 µg gH/gL or 1 µg gH/gL/gp42 nanoparticles plus 1 µg of naked ferritin nanoparticle vaccine were given intramuscularly to each mouse. The bivalent formulation comprised 1 µg gH/gL or 1 µg gH/gL/gp42 nanoparticle plus 1 µg gp350D_123_ nanoparticle vaccine. Immunizations were given at weeks 0 and 3. Sera were collected -2 days before immunization, and then at week 2, 5, and 8 post-immunizations. The animal studies with gH/gL and gH/gL/gp42 were performed in separate, independent experiments. Given the limited amount of sera available from these mice, it was technically not possible to perform neutralization assays for a head-to-head comparison.

Ferrets and NHP studies were carried out in accordance with the recommendations of the Association for Assessment and Accreditation of Laboratory Animal Care International Standards and with the recommendations in the Guide for the Care and Use of Laboratory Animals of the United States—National Institutes of Health. The Institutional Animal Use and Care Committee of BIOQUAL approved these experiments. Ferrets (n=6/group) were injected intramuscularly with 15 µg gH/gL or 15 µg gH/gL/gp42 nanoparticles plus 15 µg gp350D_123_ nanoparticle vaccine in the presence of Sanofi Pasteur AF03 adjuvant at 50% (v/v) formulation at weeks 0 and 4. Sera were analyzed at weeks 0, 2, and 6. All rhesus macaques are considered rhLCV-seropositive as pre-immune sera showed high background against gH/gL and gH/gL/gp42 and residual activity against gp350 (Supplemental Figure 6). Rhesus macaques (n=4/group) were injected intramuscularly with 25 µg gH/gL or 25 µg gH/gL/gp42 nanoparticles plus 25 µg dose of gp350D_123_ nanoparticle vaccine in the presence of Sanofi Pasteur AF03 adjuvant at 50% (v/v) formulation at weeks 0, 4, and 10. Sera were analyzed at weeks -1, 2, 6, 8, and 12.

### Enzyme-linked immunosorbent assay

Plates were coated with antigens at 100 ng/well in PBS and incubated at 4°C overnight. The plates were then washed five times in PBS-T and blocked with buffer containing 5% milk (Difco #232100) and 1% BSA (Sigma #A906-500G) in PBS-T (BioVision #2310-100). Serial dilutions of serum were made in 2.5% milk and 0.5% BSA in PBS-T. The diluted sera were added to the plate and incubated for 1 hour at room temp before being washed five times in PBS-T. Anti-mouse-HRP secondary (GE NA931V), anti-NHP-HRP secondary (Invitrogen), or HRP anti-ferret IgG (LS Bio LS-C61236-1) were added to the plate, and incubated for 1 hour at room temperature. The plate was washed five times and Sure Blue Substrate (KPL #52-00-00) was added at 100 µL/well. Once color was visualized, the reaction was stopped by adding 100 µL of 1N H_2_SO_4_ and a Spectramax M5 plate reader was used to measure absorbency at 450nm.

### GFP reporter virus neutralization assay

Immune sera from vaccinated mice, ferrets, or monkeys were serial diluted and incubated with B95-8/F EBV GFP-reporter virus for 2 hours. The mixture was added to Raji B cells, SVK CR2 or 293 epithelial cells and incubated for 3 days (*15*). Cells were then washed and fixed for flow cytometry to measure for GFP-positive cells

### Quantification of antibody titers in plasma by luciferase immunoprecipitation system assay

EBV gp350, gH/gL, and gp42 antibody titers in the week 1 post EBV challenge plasma samples were measured by luciferase immunoprecipitation system (LIPS) assay as previously described (*35, 36*). Briefly, cell lysates expressing EBV gp350, gH/gL, or gp42 Renilla luciferase fusion proteins were incubated with plasma from week 1 post infection for 1 hr and immunoprecipitated with protein A/G beads for 1 hr. Coelenterazine substrate was added to each well and luciferase activity was measured in light units (LU) by a luminometer. Each sample was tested in duplicate. The inoculum for the passive transfer in vivo study, purified IgG from single chain gH/gL-NP+gp350D_123_-NP or single chain gH/gL/gp42-NP+gp350D_123_-NP immunized BALB/c mice, was serially diluted and used to generate a standard curve. Plasma from mice 1 week post EBV challenge that received purified IgG from naïve BALB/C mice were included in each plate as a negative control. LU values that were in the linear range of a standard curve were converted to antibody titers in the plasma by interpolating a standard curve using GraphPad PRISM software.

### Passive transfer EBV challenge study

Humanized mouse experiments were carried out in accordance with federal regulations and NIH guidelines and were approved by the Animal Care and Use Committee of the National Institute of Allergy and Infectious Diseases. Three groups of BALB/c mice (n=80/group) were immunized with 5 µg gH/gL or 5 µg gH/gL/gp42 nanoparticle plus 5 µg dose of gp350D_123_ in the presence of AF03 adjuvant at weeks 0, 3 and 7 to induce an antibody response. Sera from each of the vaccinated BALB/C groups were pooled at weeks 4, 5, 8, 9, and 10 for mouse IgG purification. The control group was unimmunized BALB/c mice; similar non-immune sera have often been used as a negative control in previous publications (*37, 38*). Purified mIgG from each group was passively transferred to CD34+ humanized NSG mice (Jackson Laboratory) intraperitoneally at 20 µg of mIgG per gram of mouse at day -1, day 0, and day 1. The mice were challenged intravenously at day 0 with 10^5^ Green Raji Units of EBV. Mice were weighed weekly. EBV viremia was measured using qPCR to detect the viral gene BamH1 W (*39*) in the blood at week 5, 7, 9, 14, and 18.

### Dynamic light scattering

DLS measurements were performed at 25°C using a DynaPro Plate Reader II (Wyatt Technology). The samples were diluted in PBS, adjusted to 0.01 mg/mL concentration for each measurement. The average particle size was quantified from ten measurements.

### Immunohistochemistry

Tissues from mice were collected 27 weeks after EBV challenge and fixed in 10% neutral-buffered formalin. Sections were stained with hematoxylin and eosin, antibody to human CD20 followed by 3,3’ diaminobenzidine as a chromogen, and in situ hybridization was performed using a riboprobe for EBV EBER1. Sections were coded and read by a pathologist in a blinded fashion.

### Statistics

*P-*values were derived by Student’s *t* test or Fisher’s exact test with GraphPad PRISM version 9.3.

## Acknowledgements

We thank all members of Sanofi Breakthrough Lab for insightful discussions throughout this study. We thank Hanne Andersen Elyard, Laurent Pessaint and Jake Yalley from Bioqual for assistance with Ferret and NHP studies. We thank Harvard Medical School Electron Microscopy Facility for negative stain transmission electron microscopy. We thank Amy Sullivan, Kelly Balko, and the Sanofi Comparative Medicine group for help with the mouse immunogenicity studies,

## Author contributions

C.-J. W., W.B., L.A.N., J.I.C., and G.J.N. designed research studies; L.A.N., W.B., J.D.B, R.K., S.P.; J.R.F., T.-H.C. performed the research; L.A.N., C.-J. W., W.B., J.D.B., R.K., S.P.; J.R.F., J.I.C., and G.J.N. interpreted and discussed the data; C.-J. W., W.B., L.A.N., J.I.C., and G.J.N. wrote the paper and all authors participated in manuscript revisions.

## Data and materials availability

All data is available in the main text or the supplementary materials.

## Supplementary Materials (on a separate file)

Materials and Methods

Figs. S1 to S9

Tables S1 to S2

---

**Table S1.**
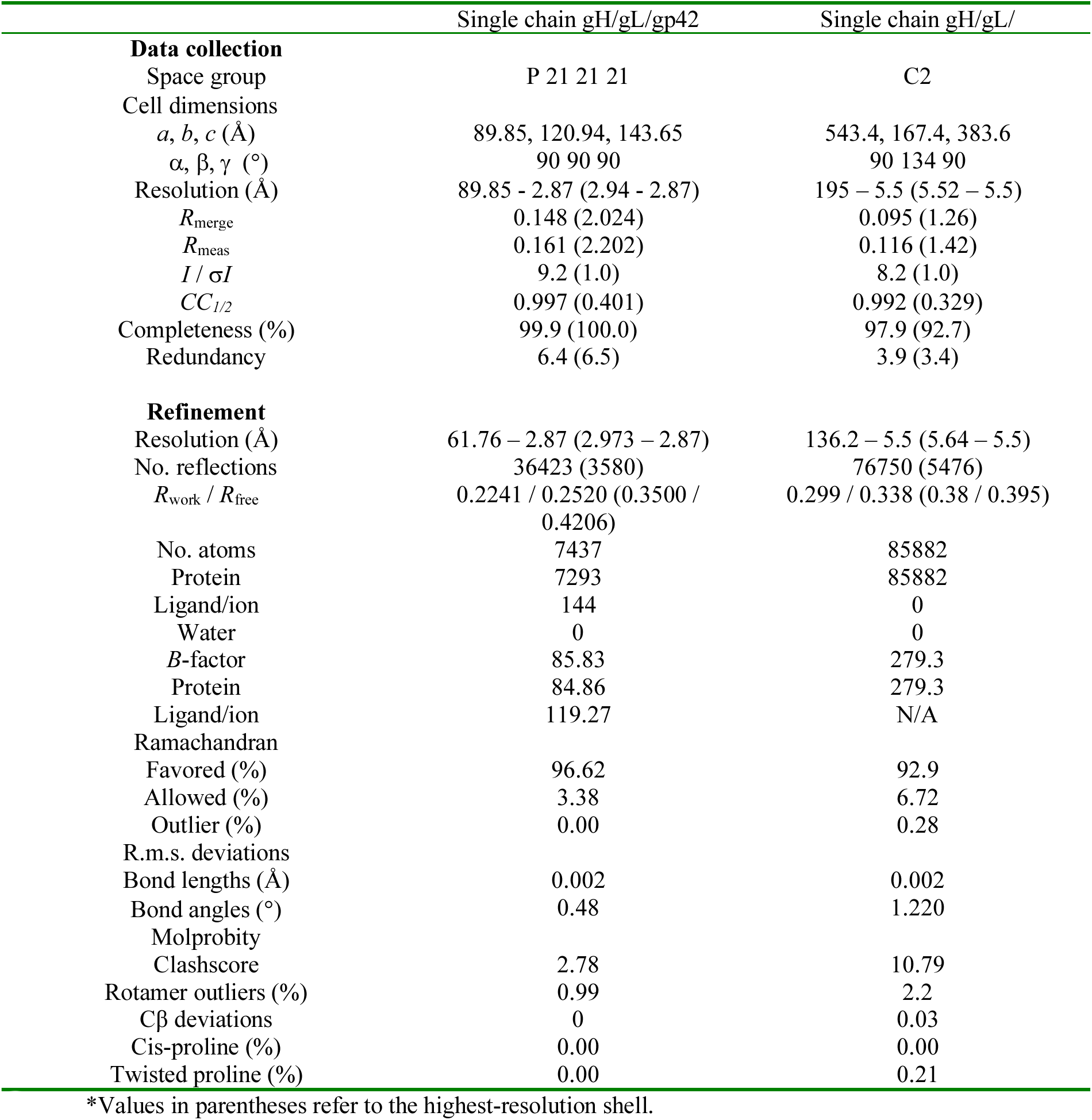
Data collection and refinement statistics (molecular replacement)

**Table S2.**
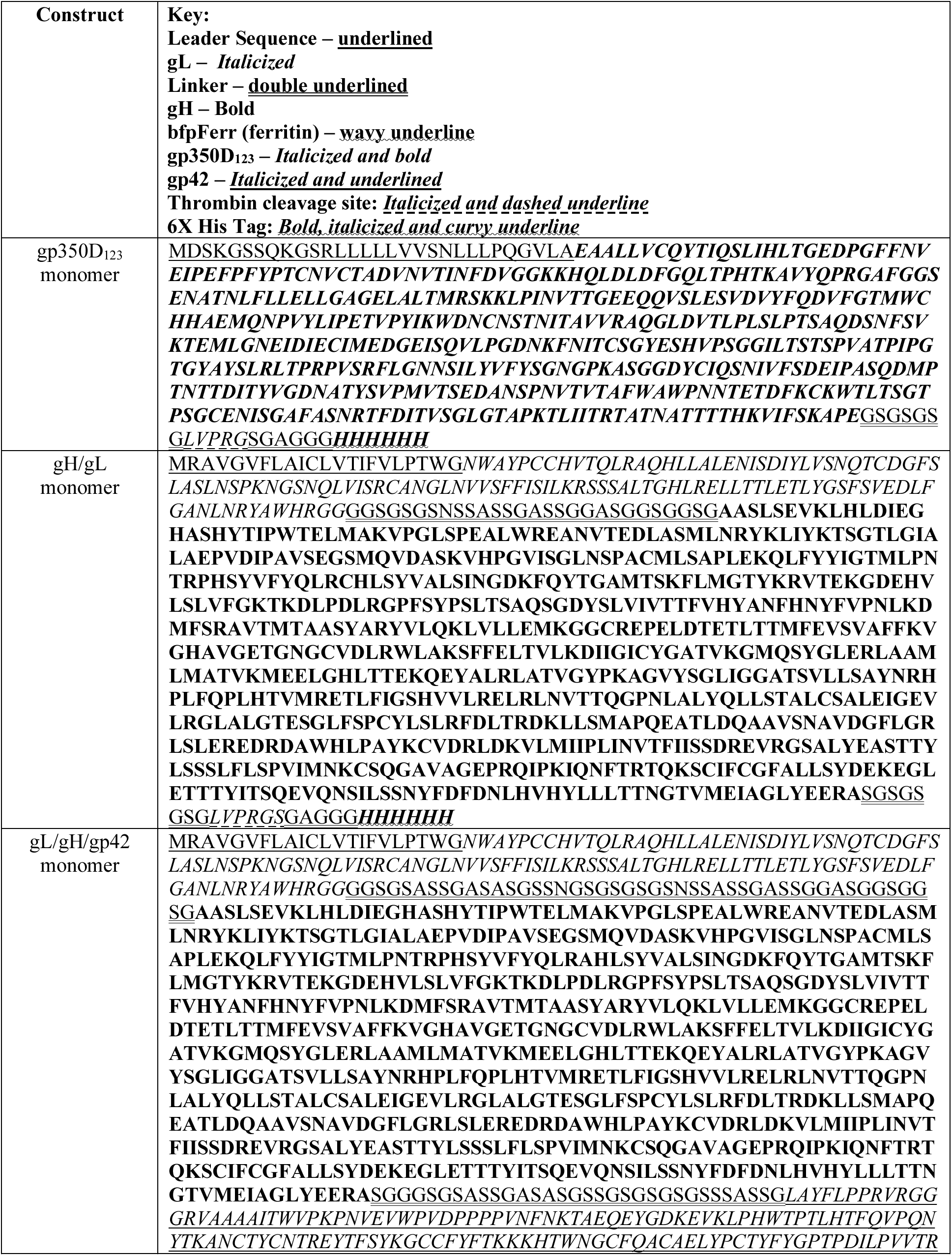
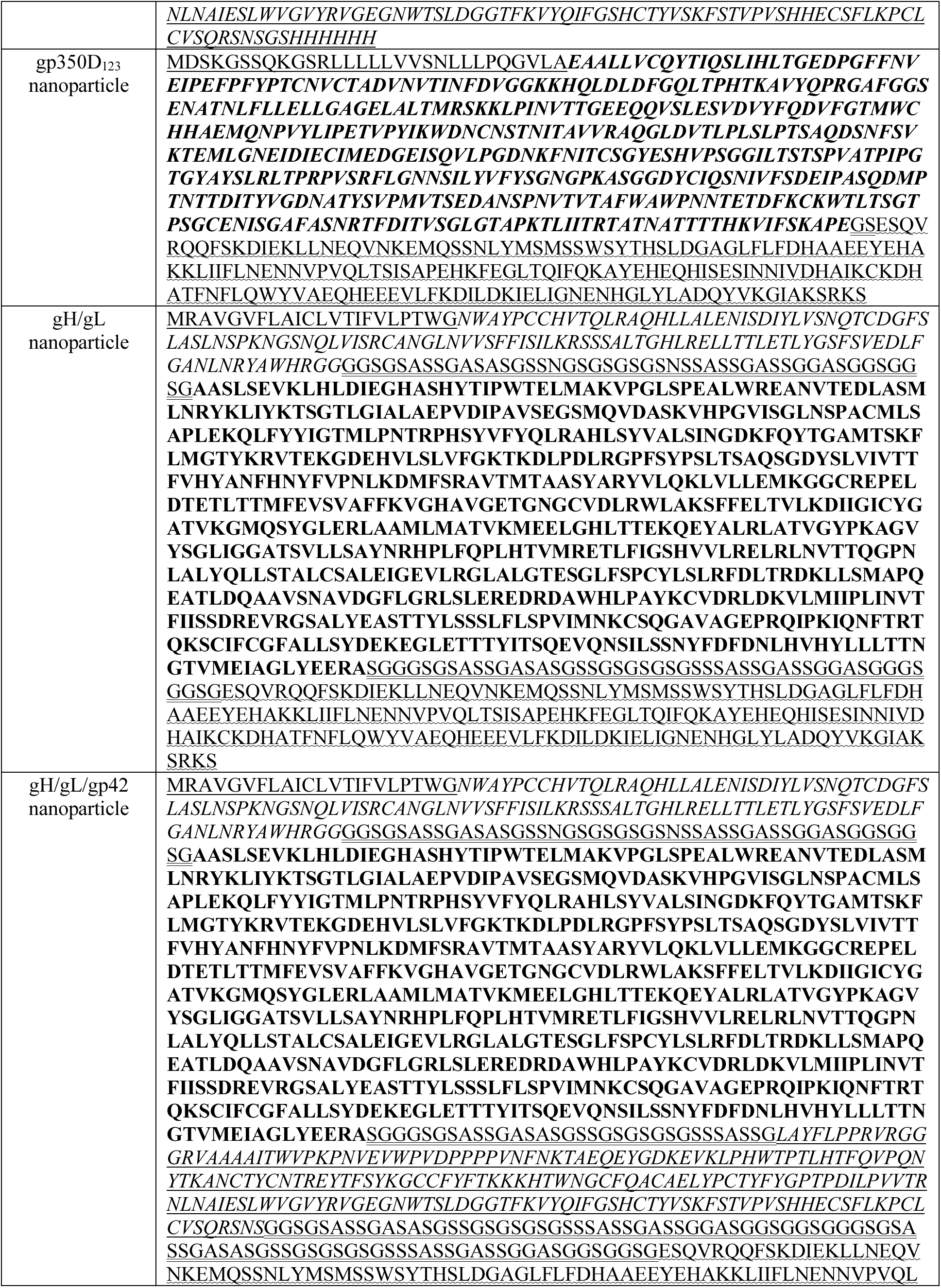
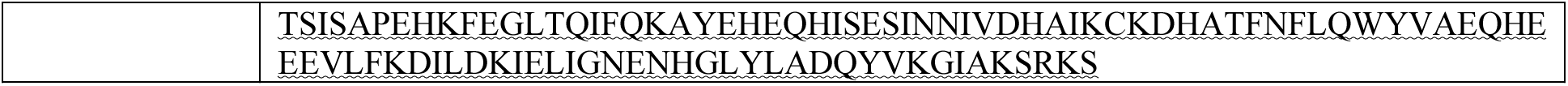
Amino acid sequences of constructs used.

## SUPPLEMENTAL FIGURES

**Supplemental Figure 1.**
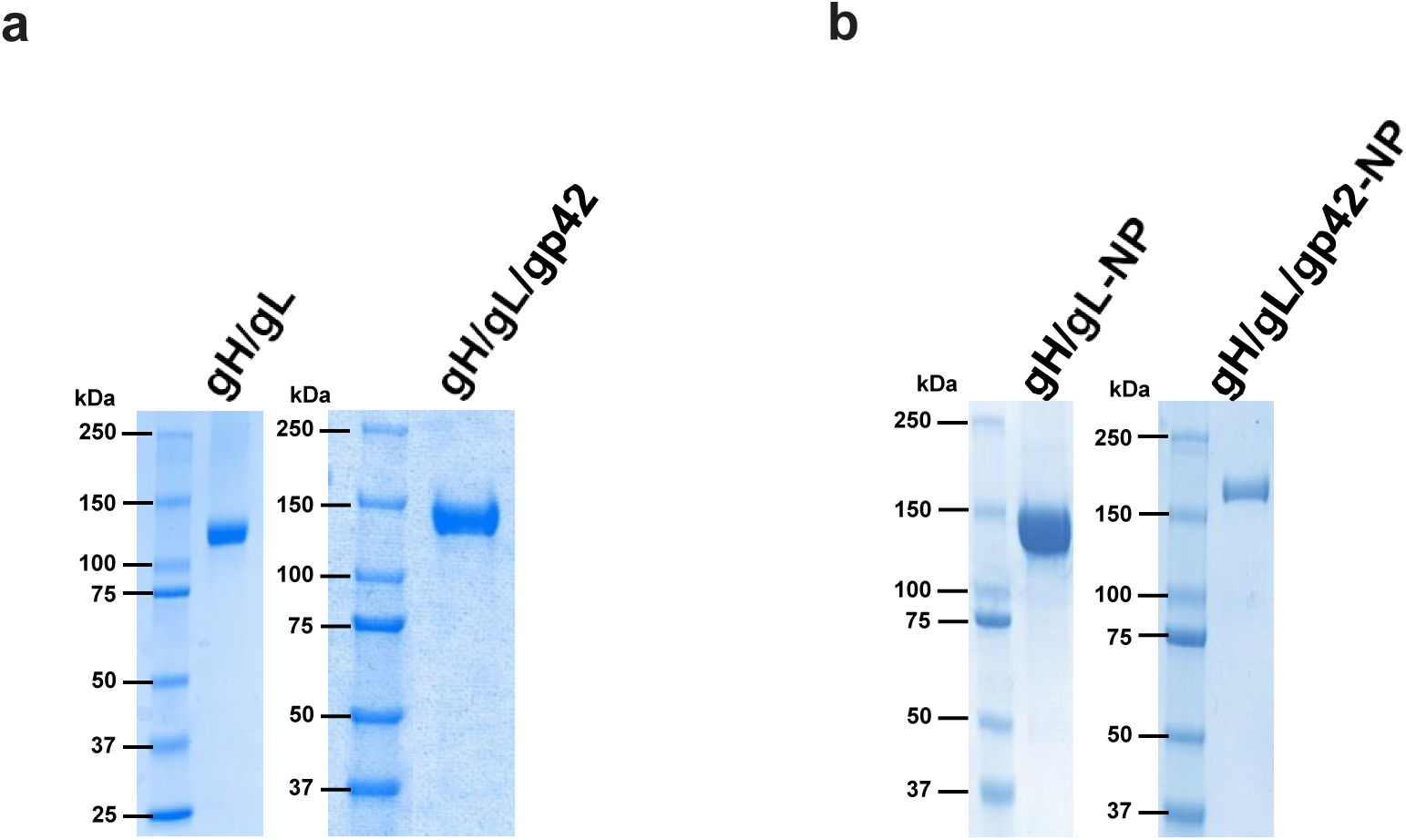
Expression of single chain gH/gL and single chain gH/gL/gp42 constructs. (A.) SDS-PAGE analysis of purified single chain gH/gL and single chain gH/gL/gp42 constructs. (B.) SDS-PAGE of purified single chain gH/gL-NP and single chain gH/gL/gp42-NP.

**Supplemental Figure 2.**
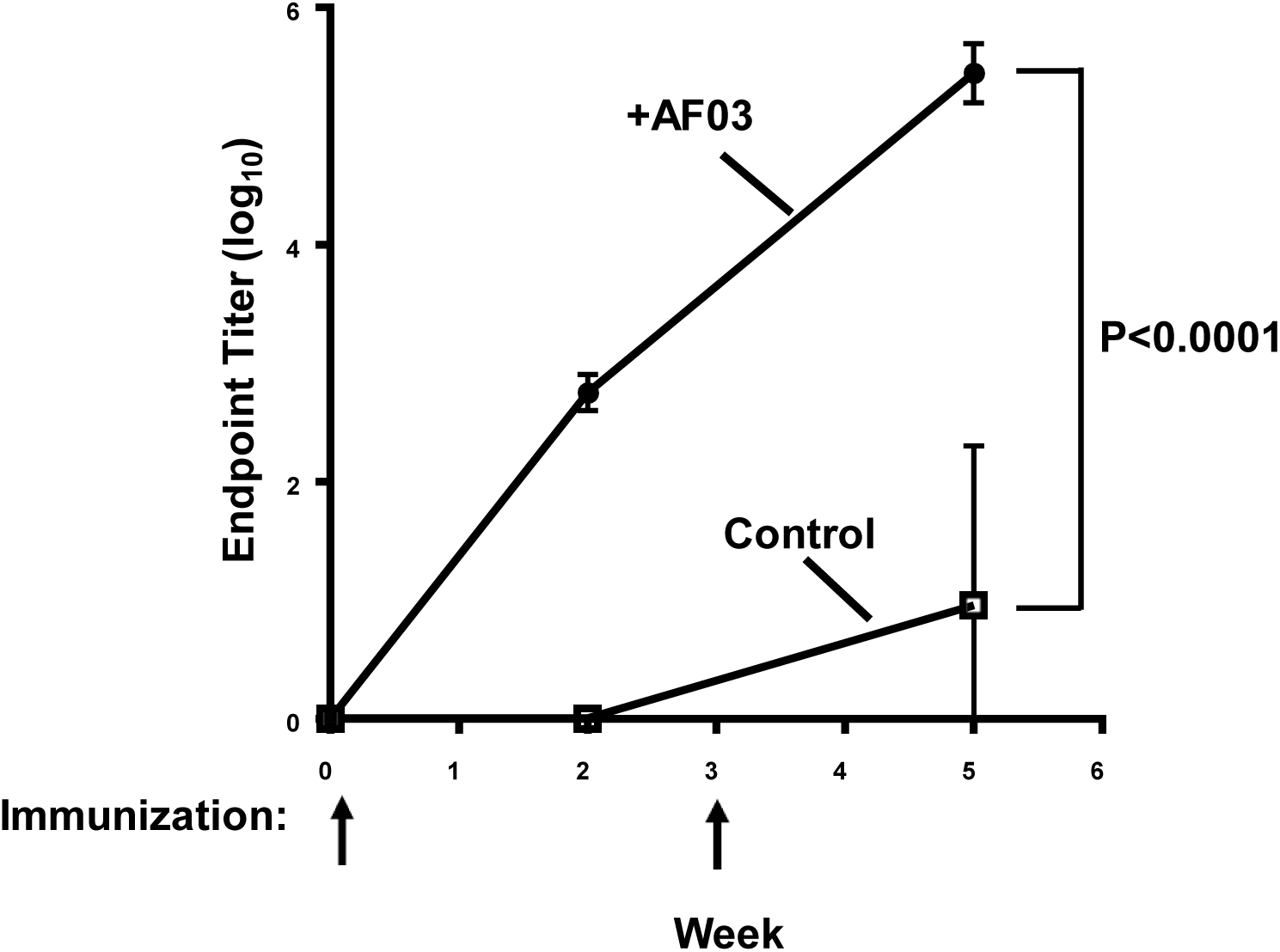
Immunogenicity of single chain gH/gL-NP with or without AF03 adjuvant. Mice were immunized at week 0 and week 3. Immune sera were collected at weeks 0, 2 and 5 and antibody titers were determined by ELISA. Mean and standard error are shown. The p value from week 5 was <0.0001.

**Supplemental Figure 3.**
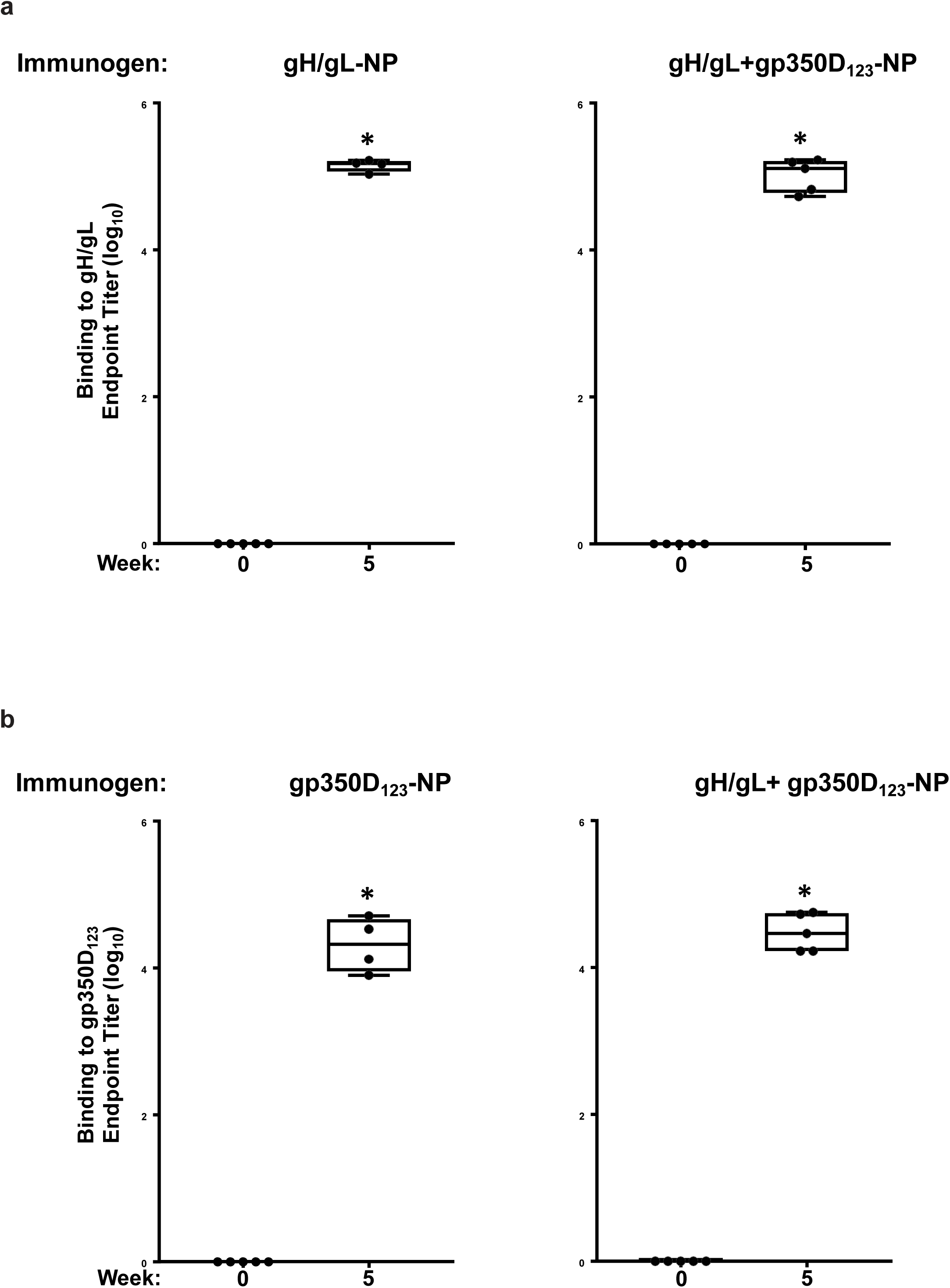
Immunogenicity of single chain gH/gL-NP and gp350D_123_-NP in mice. A monovalent single chain gH/gL-NP or gp350D_123_-NP, or bivalent single chain gH/gL-NP+gp350D_123_-NP was used to immunize mice with AF03 as adjuvant. Antibody titers pre- and post-immunization against either (a) gH/gL heterodimer or (b) gp350D_123_ were determined by ELISA. The data are shown as box-and-whiskers plots (box indicates lower and upper quartiles with line at median, and whiskers span minimum and maximum data points; *p<0.0001 compared to pre-immune sera).

**Supplemental Figure 4.**
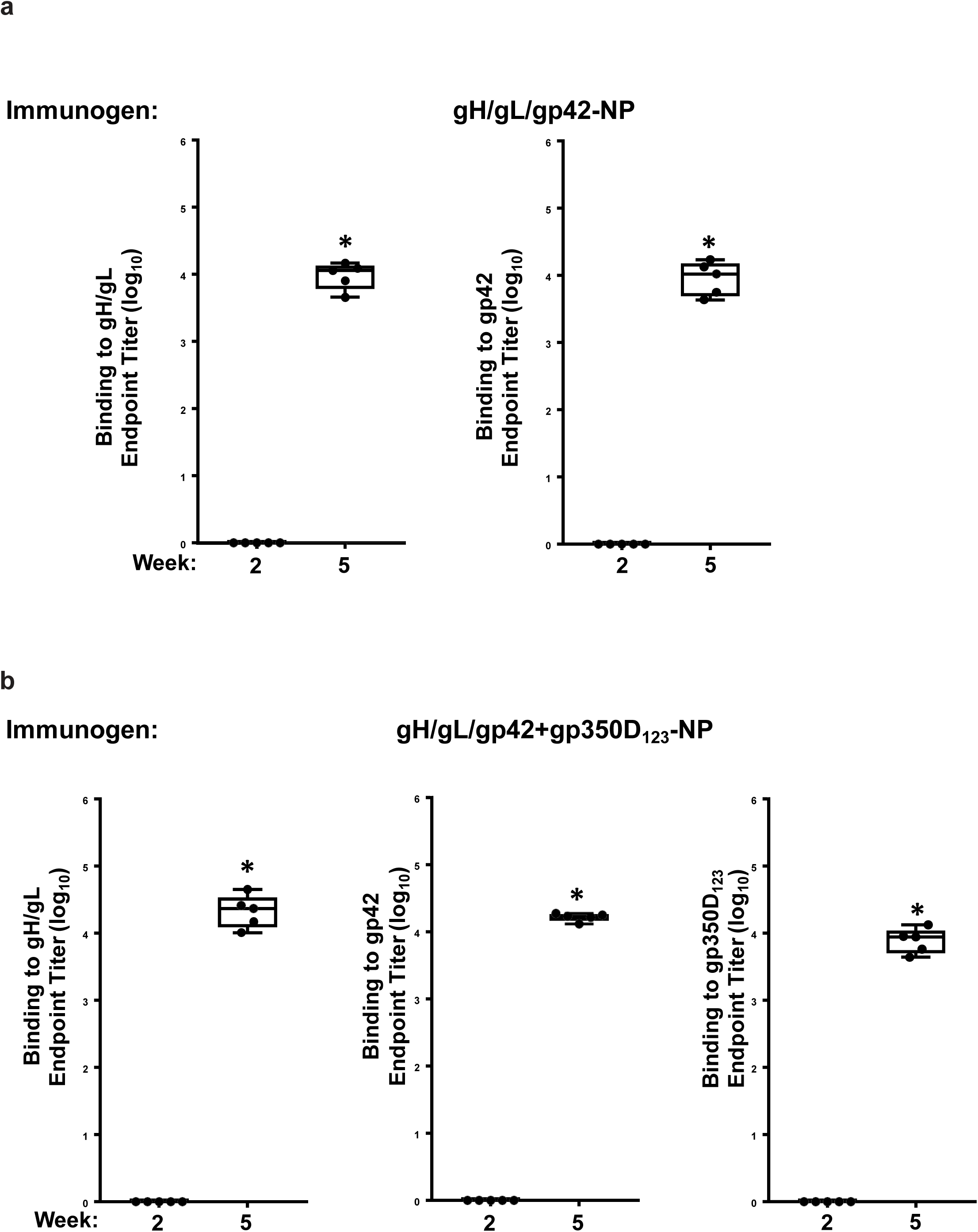
Immunogenicity of single chain gH/gL/gp42-NP with gp350D_123_-NP in mice. Monovalent single chain gH/gLgp42-NP (a) or bivalent single chain gH/gL-NP/gp42-NP+gp350D_123_-NP (b) was used to immunize mice with AF03 as adjuvant. Antibody titers from immune sera collected 2 weeks after first and second immunizations against either gH/gL heterodimer, gp42 or gp350D_123_ were determined by ELISA. The data are shown as box-and-whiskers plots (box indicates lower and upper quartiles with line at median, and whiskers span minimum and maximum data points; *p<0.0001 compared to week 2 sera).

**Supplemental Figure 5.**
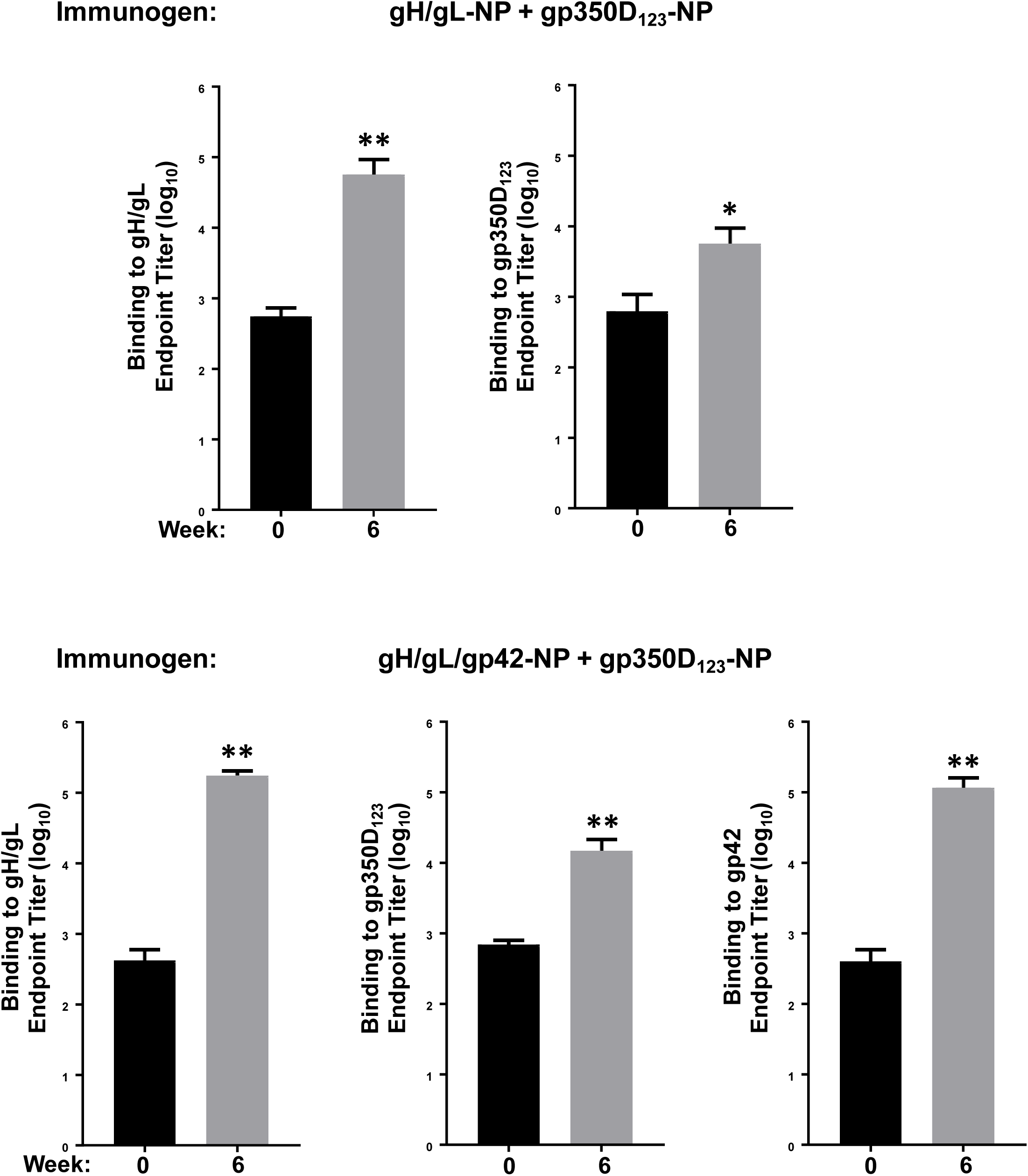
Immunogenicity of single chain gH/gL-NP+gp350D_123_-NP and single chain gH/gL/gp42-NP+gp350D_123_-NP vaccines in ferrets. Ferrets (n=6/group) were immunized with either (a) bivalent gH/gL-NP+gp350D_123_-NP or (b) bivalent gH/gL/gp42-NP+gp350D_123_-NP vaccines at weeks 0 and 4. Binding antibody titers to gH/gL, gp350D_123_, and gp42 were determined. Means and standard deviations are shown (**p<0.0001 or *p<0.05 respectively compared to pre-immune sera).

**Supplemental Figure 6.**
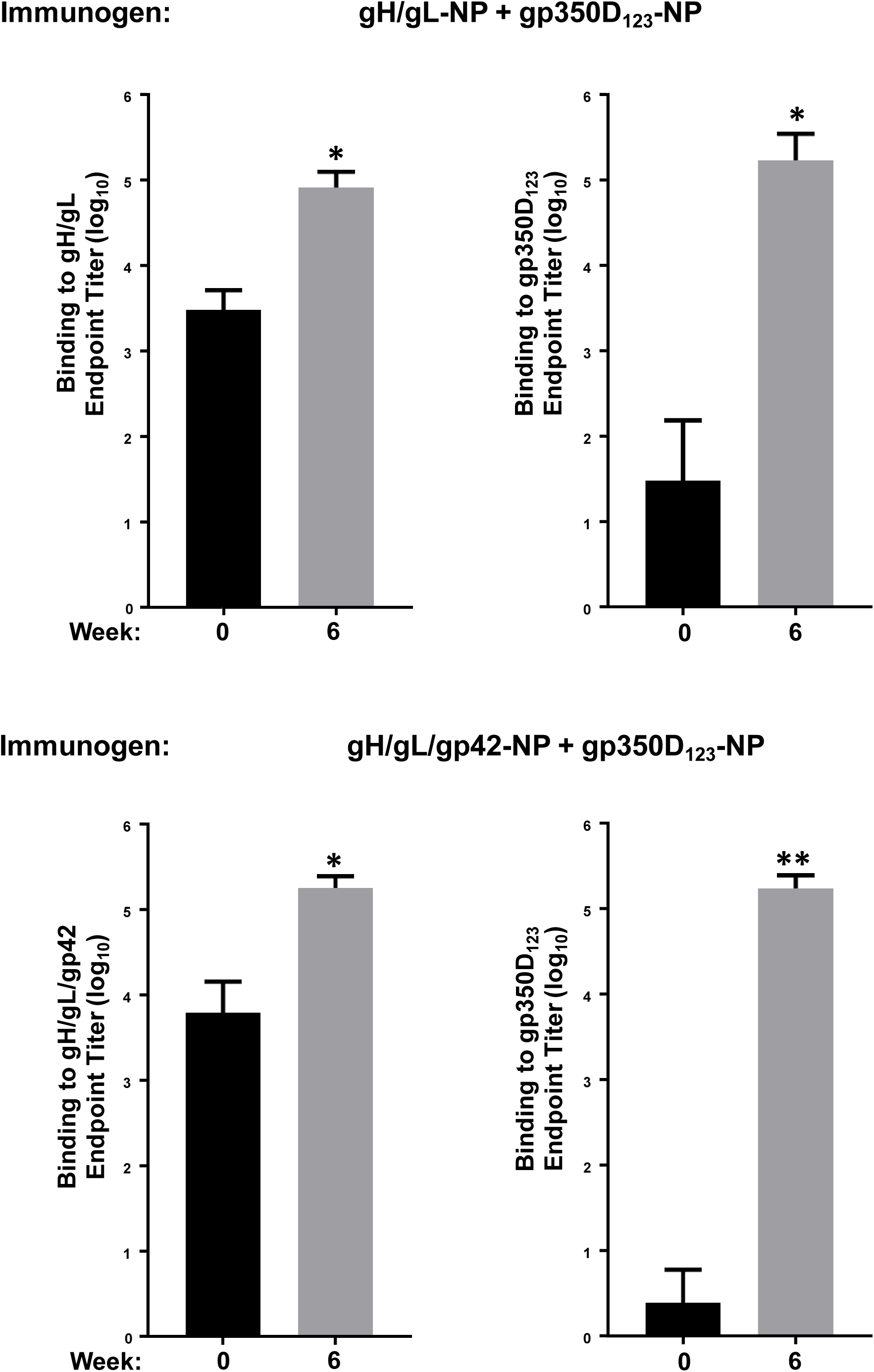
Immunogenicity of single chain gH/gL-NP+gp350D_123_-NP and single chain gH/gL/gp42-NP+gp350D_123_-NP vaccines in nonhuman primates. Rhesus macaques were immunized with either (a) single chain gH/gL-NP+gp350D_123_-NP or (b) single chain gH/gL/gp42-NP+gp350D_123_-NP bivalent vaccines at weeks 0, 4, and 10. Pre- and post-immunization binding antibody titers against gH/gL or gH/gL/gp42 or gp350D_123_ were determined by ELISA. Means and standard deviations are shown (**p<0.0001 or *p<0.01 respectively compared to pre-immune sera).

**Supplemental Figure 7.**
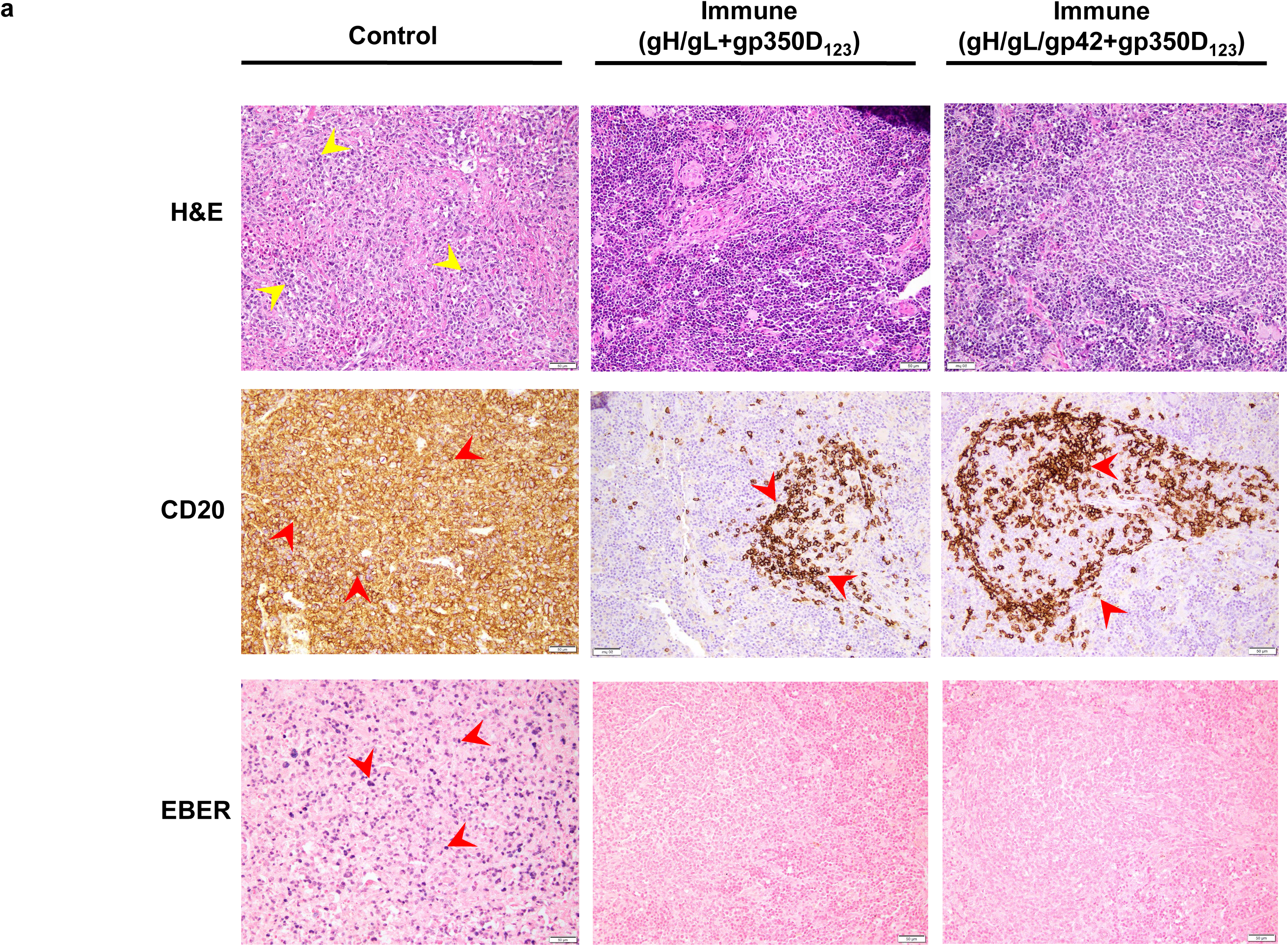
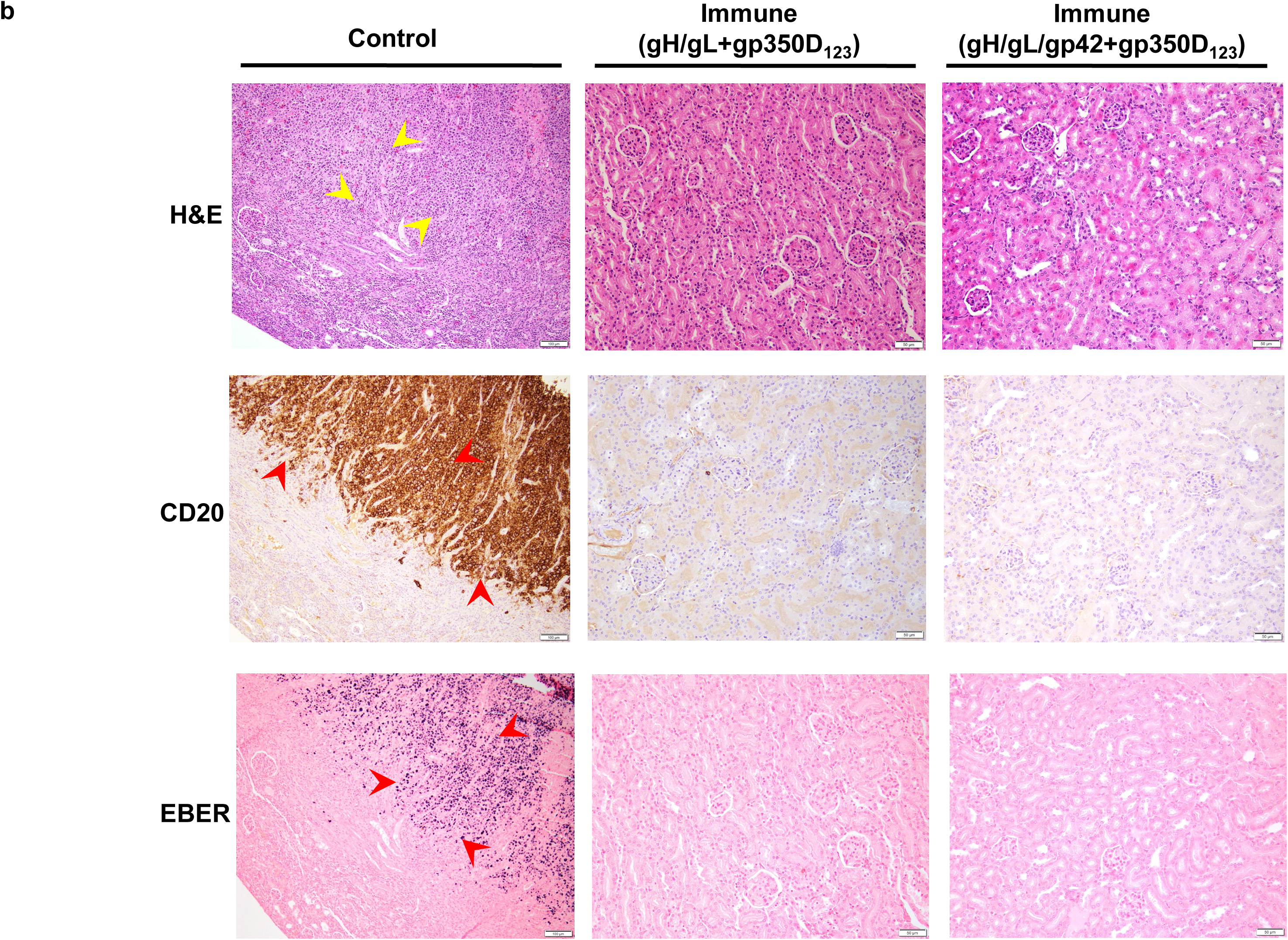
EBV lymphoma in challenged humanized mice. EBV-positive B cell lymphomas and CD20 and EBER positive cells were observed in the (a) spleen and (b) kidney of mice receiving IgG from naïve BALB/C after EBV challenge (Control). No evidence of EBV-positive B cell lymphomas was seen in mice receiving IgG from animals vaccinated with single chain gH/gL-NP+gp350D_123_-NP or gH/gL/gp42-NP+gp350D_123_-NP (Immune). CD20 staining and EBER are also negative in these mice. Tissues were harvested and stained as in Fig. 5. Arrows indicate representative areas of pathology and staining for the indicated cellular or viral proteins.

**Supplemental Figure 8.**
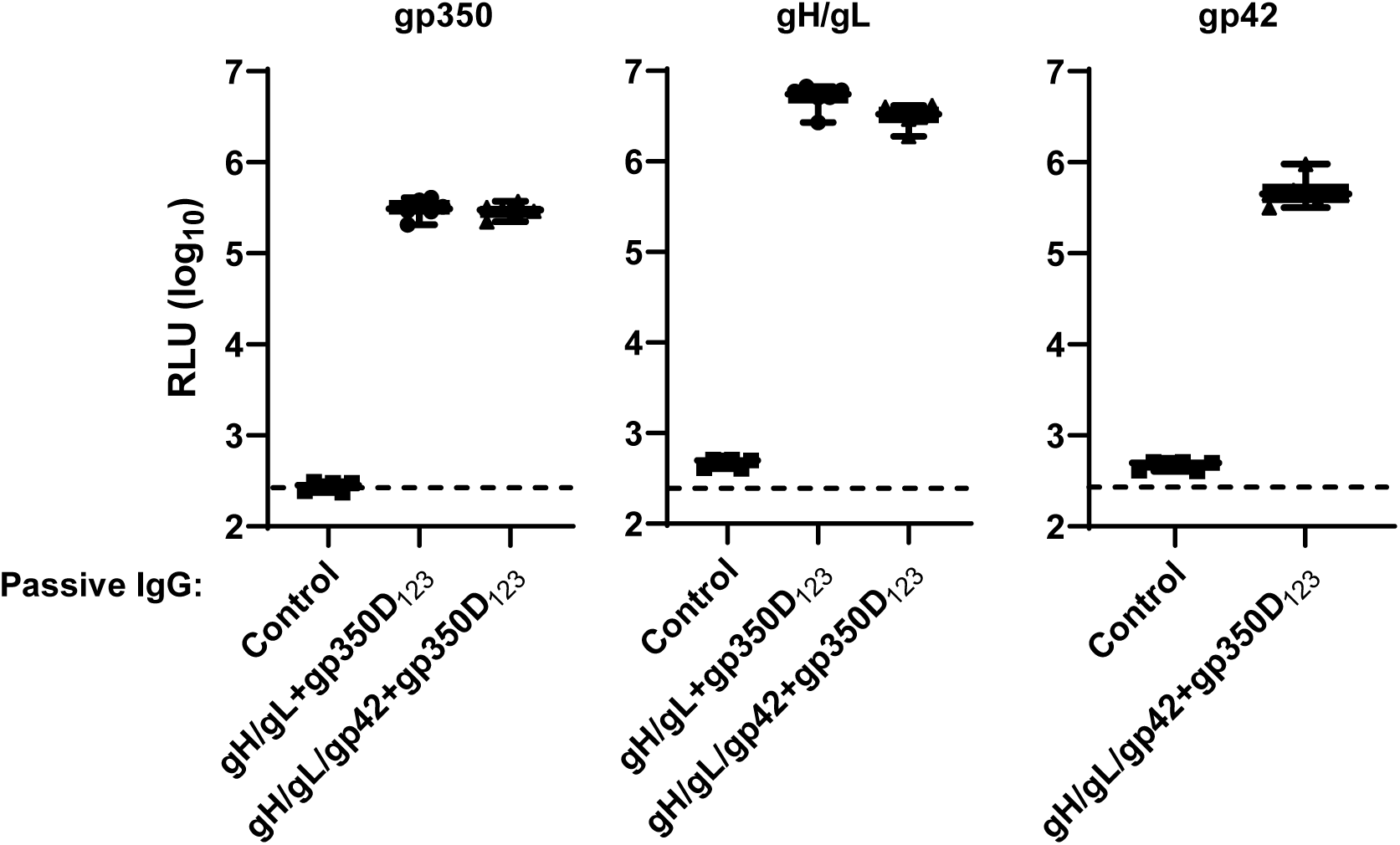
Anti-gH, gL, and gp42 antibody levels in mice challenged with EBV. Antibody titers in plasma samples of humanized mice receiving IgG from vaccinated or naïve (control) mice obtained one week after challenge were measured by LIPS assay and shown as RLUs. Antibody titers in animals that received IgG from naïve, single chain gH/gL-NP+gp350D123-NP, and gH/gL/gp42-NP+gp350D123-NP immunized BALB/c mice are shown. Each symbol indicates one mouse. Minimum and maximum data points are represented by whiskers and box represents upper and lower quartiles with the horizontal line at the median.

